# Embracing naturalistic paradigms: substituting GPT predictions for human judgments

**DOI:** 10.1101/2024.06.17.599327

**Authors:** Xuan Yang, Christian O’Reilly, Svetlana V. Shinkareva

## Abstract

Naturalistic paradigms can assure ecological validity and yield novel insights in psychology and neuroscience. However, using behavioral experiments to obtain the human ratings necessary to analyze data collected with these paradigms is usually costly and time-consuming. Large language models like GPT have great potential for predicting human-like behavioral judgments. The current study evaluates the performance of GPT as a substitute for human judgments for affective dynamics in narratives. Our results revealed that GPT’s inference of hedonic valence dynamics is highly correlated with human affective perception. Moreover, the inferred neural activity based on GPT-derived valence ratings is similar to inferred neural activity based on human judgments, suggesting the potential of using GPT’s prediction as a reliable substitute for human judgments.

## I. INTRODUCTION

TRADITIONAL psychology research has predominantly relied on highly controlled laboratory tasks, typically employing simplistic and carefully manipulated stimuli cues [1]. The ecological validity of this approach is questionable, a situation leaving a gap in our understanding of the complicated psychological and neural dynamics present in the ongoing, hierarchical, and multimodal processing associated with real-life contexts. To address this issue, naturalistic paradigms have been used increasingly often in research. These paradigms employ rich, continuous, and multimodal stimuli approximating some of our daily experiences, such as watching movies, listening to stories, and interacting with people. These approaches have a higher ecological validity and reproducibility and can yield novel insights [1], [2], [3].

However, methodological obstacles associated with extracting the features required for analyzing naturalistic paradigms in an accurate, unbiased, and cost-efficient manner prevent researchers from widely embracing this approach. Features are typically obtained through behavioral experiments, such as moment-by-moment affective ratings of narratives, annotation of discrete emotions in movie scenes, or specific behaviors during interactions. Obtaining these features through subjective ratings is usually time-consuming and costly. Moreover, they can be biased by the intermittent nature of the rating process, stimuli repetition, and social desirability.

With the rapid advancement of artificial intelligence, large language models (LLMs) such as GPT have demonstrated great potential for predicting human-like behavioral judgments on concepts traditionally deemed inherently human, including morality [4], decision-making [5], and social desirability [6]. Although these investigations have used simple and carefully phrased instructions for GPT’s predictions, the most recent release of the GPT model can process longer text and understand the requirement of multifaceted tasks. This advancement lays the groundwork for employing LLMs to automate behavioral ratings of complex naturalistic stimuli. Moreover, provided that the pre-trained LLMs faithfully capture average human judgments, the reproducible ratings produced by time-efficient, cost-effective, and universally accessible LLMs would be invaluable for automating the feature extraction process, and, thus, could eliminate the methodological barrier in the application of naturalistic paradigms.

The current study evaluates the feasibility of using GPT as a substitute for human ratings of hedonic valence in narratives. We instructed human participants and GPT to rate valence for four narratives. We also extracted the lexical-level valence values from the psycholinguistic metabase (SCOPE) [7] to examine if GPT valence ratings of narrative segments outperform ratings estimated from the superimposition of the hedonic valence associated with individual words taken out of their context. The human ratings, GPT ratings, and SCOPE values were correlated with fMRI data collected in a separate experiment. The participants used for human ratings and neuroimaging were different but they listened to the same narratives. Using affective ratings obtained from a different group of participants than the sample used for fMRI or EEG recording has been shown to produce accurate results and is a well-established practice [8], [9], [10].

## II. METHODS

### A. Narrative selection

To illustrate the potential of using GPT ratings as an alternative to human ratings for affective neuroscience, we leveraged narrative stimuli from the publicly available Narrative fMRI dataset [11]. Four narratives (“I Knew You Were Black”, “The Man Who Forgot Ray Bradbury”, “Slumlord”, and “Reach for the Stars One Small Step at a Time”) were selected based on the narrative length (14.02 minutes and 2256 words on average) to fit a one-hour human behavioral experiment.

### B. Stimuli segmentation

We parsed narrative transcripts into segments to effectively capture affective fluctuations within sentences. Narratives were parsed into an average of 226.75 standalone segments, with every segment lasting 3.26 seconds on average and capturing phrases, generally adhering to grammatical rules and natural speech pauses. Two research assistants parsed the transcripts independently and reached a consensus. Written transcripts were used for the behavioral rating experiments.

### C. Human behavioral experiments

To obtain human valence ratings, we conducted four online experiments on the SONA system (https://www.sona-systems.com) under a study protocol approved by the Institutional Review Boards (IRBs) of the University of South Carolina. Undergraduate students (*N* = 176) participated in these experiments. In each experiment, participants read the narrative transcript one segment at a time and reported their feelings using a valence-by-arousal grid (see the *Appendix A* for detailed instructions). Participants could click anywhere on the grid, resulting in a continuous measure for valence (range from -4.5 to 4.5, with -4.5 representing very negative and 4.5 representing very positive valence) and arousal (range from 0 to 9, with 0 representing low arousal and 9 representing high arousal). At the beginning of each experiment, two paragraphs with positive and negative valence, respectively, were displayed to illustrate the extremes of valence content present in the experiment. Participants were asked to calibrate their scoring to utilize the full valence and arousal scales. To confirm participant engagement, comprehension questions were asked at the end of each experiment. Nineteen participants were excluded due to low comprehension accuracy (less than 60%), resulting in a final sample of 157 participants across the four experiments (*M*_age_ = 20.75, *SD*_age_ = 4.01, 132 females). Previous research suggests that a sample size of over 25 participants yields consistent valence ratings [12]. To decrease the effect of idiosyncratic affective ratings in some participants, we used the STATIS procedure [13] to downweight the contribution of participants who share low-scoring similarity with the sample. The weighted average ratings were rescaled using min-max scaling to match the scales of GPT valence ratings (from -4 to 4) and arousal ratings (from 1 to 9). The rescaled weighted average ratings were then used as the final gold standard to compare with GPT and lexical-level values.

### D. GPT queries

To evaluate the feasibility of substituting human behavioral ratings with GPT predictions, we used the same instructions (422 words in total) as those used in human behavioral experiments, with minimal modification to ensure formatted output (see the *Appendix B* for details). Considering that GPT cannot use the valence-by-arousal rating grid and can only provide integer ratings, we requested GPT to use a 9-point discrete scale to rate valence (from -4 to 4) and arousal (from 1 to 9). To ensure direct comparison, we rescaled the human ratings and the lexical-level SCOPE values to match GPT’s ratings. Instead of presenting GPT with the narrative segments one by one and making queries conversationally, we fed in both the instructions and the entire narrative to GPT in a single query. For each of the four narratives, we repeated the query 10 times using the GPT-4-preview-1106 model and all the default parameters except for the temperature. We set the temperature parameter to 0.2 and the random seed to 123. We experimented with different temperature settings and found 0.2 to provide optimal results. For each narrative, ratings across the 10 iterations were averaged and used as the final GPT ratings.

### E. Improving the performance of GPT

We used several prompting strategies to ensure that GPT could follow our instructions and to detect potential errors when GPT failed to follow the instructions. First, we defined the role of GPT as “an undergraduate student who is a native speaker of English” through the system message, which matches the demographic characteristics of our human sample. Second, to improve GPT’s understanding of the expected outcome, we adopted a Few-Shot strategy by providing the identical instructions used for human ratings in a user message and specifying the expected responses to three example segments in an assistant message. Third, in a following user message, we instructed GPT on the total number of segments present in the input prompt and fed in the entire narrative segments altogether with five hash characters (#####) inserted between segments to facilitate segment extraction by GPT. To assure alignment (i.e., that GPT valence and arousal predictions are locked to the input segment) and to detect potential misalignments (i.e., cases where GPT valence and arousal predictions are based on misaligned or hallucinated segments), we requested GPT to export the segment ID, along with the first and the last words of the segment being rated, and the valence and arousal ratings. When the exported first and last words were not aligned with our input, we inferred that GPT ratings were misaligned.

### F. Lexical-level valence and arousal values

The lexical-level valence and arousal values were used as a baseline to examine how GPT affective prediction of narrative segments may differ from the superimposition of the affective quality of single words. To maximize the number of narrative words that had available values, we leveraged three major databases (Warriner, Glasgow, and NRC) from the SCOPE Metabase [7]. These databases include measures on valence and arousal through normative behavioral experiments (e.g., subjectively rate the valence and arousal on individual words, or make an affective judgment on which word is more positive/negative compared to other presented words). To match words independently of their form, we used the NLTK Python package to lemmatize the words from the databases and the narratives before matching them. Words from the narratives that could not be matched were treated as missing. To combine the information from these three databases, we standardized the values within each database by subtracting the mean and dividing by the standard deviation of the database values. Values for words present in more than one database were averaged. We then rescaled these lexical-level averaged standard scores to match the scales of GPT valence ratings (-4 to 4) and arousal ratings (1 to 9). The rescaled averaged standard values for each word were then used as a baseline for subsequent comparisons and analyses. Across the four narratives, 59.80% of unique words (1080 out of 1806) had available valence and arousal values.

### G. Neuroimaging data

To illustrate the application of GPT ratings in an affective neuroscience study, we leveraged the functional MRI data from the Narrative dataset [11]. Specifically, 46 participants passively listened to “I Knew You Were Black” and “The Man Who Forgot Ray Bradbury” in two separate runs, and 18 participants listened to “Slumlord” and “Reach for the Stars One Small Step at a Time” in one run. Eight participants (16 scans) were excluded because of low accuracy on comprehension questions, resulting in a total sample size of 112 scans from 56 people for the current study (*M*_age_ = 22.70, *SD*_age_ = 6.65, 30 females).

We used the preprocessed data available in the Narrative dataset. This preprocessing included temporal and spatial corrections of the fMRI volumes, followed by normalization to the MNI space using fmriprep. The data were then smoothed using a 6mm FWHM Gaussian filter using 3dBlurToFWHm program in AFNI. Confounding variables, including head motions, signals in CSF and white matter, and time drifts (for details, see [11]) were regressed out using 3dTproject program in AFNI. The residuals were used as input for subsequent analyses. Finally, all images were resliced to a matching spatial resolution of 3 × 3 × 4 *mm*^3^ voxels using the 3dResample program in AFNI.

## III. ANALYSIS

### A. Simulation test

To evaluate how many human raters are needed to outperform GPT ratings, we randomly split our set of *X* = 40 human ratings per story into sets of *N* test ratings and *X* − *N* gold standard ratings, with *N* varying between 1 and 20. Then, we compared the average rating for the gold standard set to ratings from 1) GPT, 2) SCOPE values, and 3) the average rating for the test set.

### B. Neuroimaging data analyses

To account for the within-subject variance associated with the same participant listening to different narratives, scans acquired in different runs were specified as multi-sessions in SPM12 (www.fil.ion.ucl.ac.uk/spm/). We conducted a parametric modulation regression to examine how the Blood Oxygen Level Dependent (BOLD) signal in different brain regions varies as a function of hedonic valence while controlling for arousal. The parametric regressor contained three vectors, one for the onset time of narrative stimuli (the beginning of the segments for Human and GPT; the beginning of the words for lexical-level), and two for the continuous valence and arousal scores obtained from the human ratings, GPT ratings, and lexical-level values. All three vectors were convolved with the canonical hemodynamic Response Function (HRF) and used as input into the linear regression model as independent variables. We conducted random effects analyses to create activation maps at the group level. A cluster-size-based correction was applied. For each model, the threshold cluster size was estimated by the Gaussian Random Field method implemented in SPM12 on the basis of the number of adjacent voxels surviving a voxel-wise threshold of *p* <.001. The largest cluster size (>22 voxels) was used for comparison across models.

## IV. RESULTS

GPT valence ratings were highly correlated with human valence ratings (mean correlation coefficient = 0.81, *ps <*.00001; Fig. 1E). GPT ratings closely reproduced the U-shape relationship between arousal and valence (Fig. 1C) and the valence dynamics across time observed in human ratings (Fig. 1D), suggesting that GPT reliably emulates human affective judgments. Critically, the correlation with human ratings was remarkably higher for GPT than lexical-level values, revealing that the prediction of valence by GPT for narrative segments is not a simple superimposition of the affective quality of single words. In addition, when presented with the entire narrative and the same complex and multifaceted instructions used in human behavioral experiments, GPT successfully identified the segments, rated them on a 9-point scale for valence and arousal, and exported the results (Fig. 1A-B). We made 40 queries, with 10 repetitions per story. the total number of output segments was largely consistent with our input. On average, only 7.54% of the segments did not match our input. Critically, valence and arousal ratings across the entire narrative were remarkably consistent with our gold standard human ratings (see Fig 1D).

**Fig. 1.**
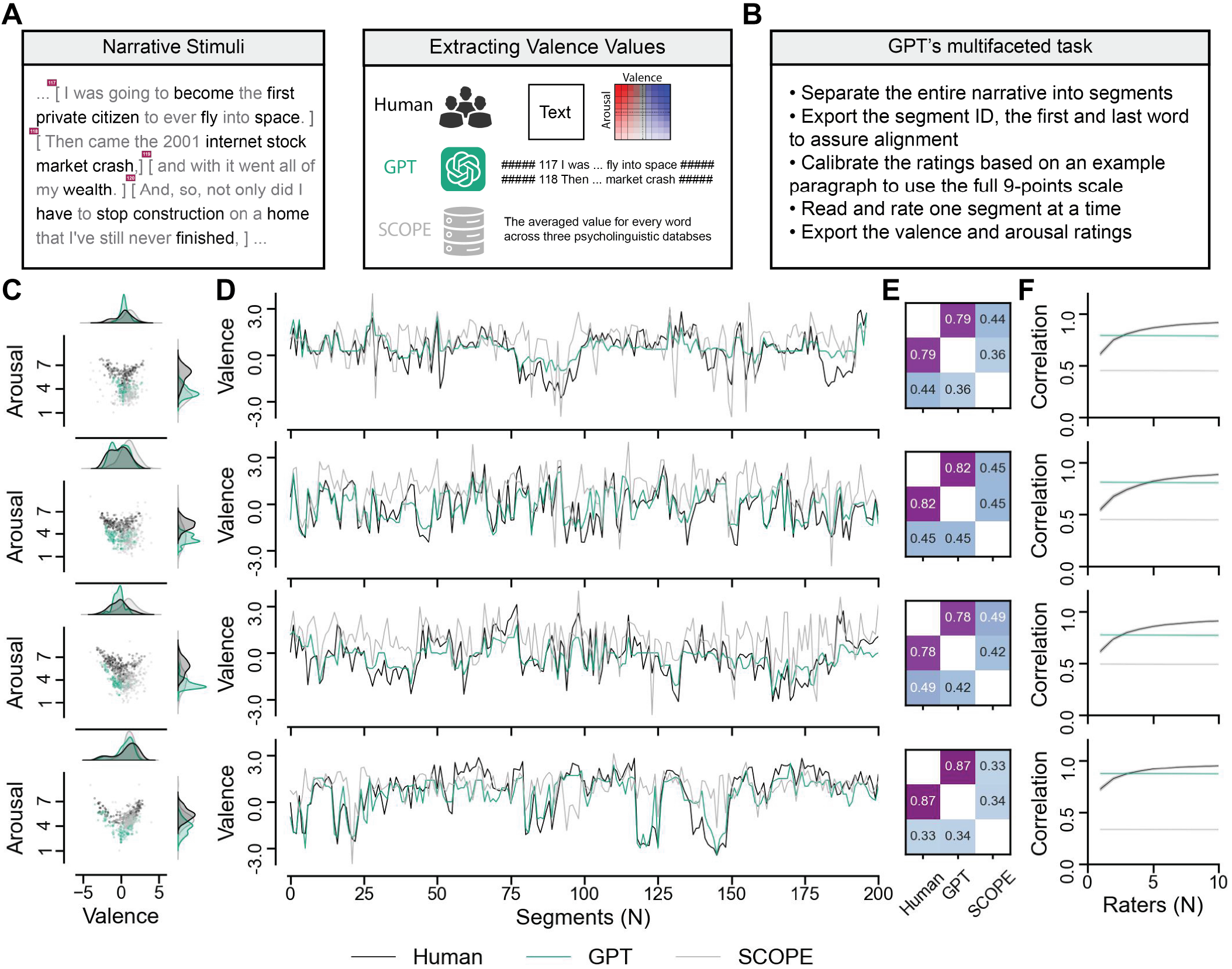
(A) Parsing the narrative transcript into segments and obtaining affective ratings from humans, GPT, and SCOPE values. (B) Description of GPT’s tasks. (C) The bivariate scatter plot of valence and arousal ratings from humans, GPT, and SCOPE values. The distribution of the ratings is shown on the top and right sides. (D) Comparison of the average valence ratings from humans, GPT, and SCOPE values as narratives unfold. (E) Pearson correlation coefficient matrix between ratings from humans, GPT, and SCOPE values. (F) Simulation of how many human raters are needed to outperform the GPT ratings. For panels C-F, the four rows correspond to the different narratives.

Next, we analyzed how many human raters are needed to outperform GPT ratings. Although GPT did not provide fully reproducible ratings across queries, its average ratings across ten repetitions of the same query outperformed the lexical-level SCOPE values (Fig. 1C-E) and single human raters (Fig. 1F). Our simulation test revealed that we need to average ratings from at least 3-5 humans, depending on the narrative, to outperform GPT ratings. The correlation between the gold standard and the human rater test set plateaued at around *N* = 10, a value significantly lower than the 25 ratings reported necessary for stable ratings in previous studies [12]. Lastly, to illustrate the substitution of human judgments by GPT predictions in affective neuroscience, we correlated the valence ratings inferred by GPT with the hemodynamic response of participants listening to the narratives in an MRI scanner. Our results (see Fig. 2) revealed a remarkable similarity between the brain regions identified by human (Table I) and GPT valence ratings (Table II). These brain regions are part of the core-affect neural network [14] and are in line with the valence-associated areas derived from affective pictures [15], affective sounds [16], and naturalistic movies [9]. In contrast, results based on the lexical-level values were less robust (Table III), possibly reflecting the importance of contextual information in affective ratings.

**Fig. 2.**
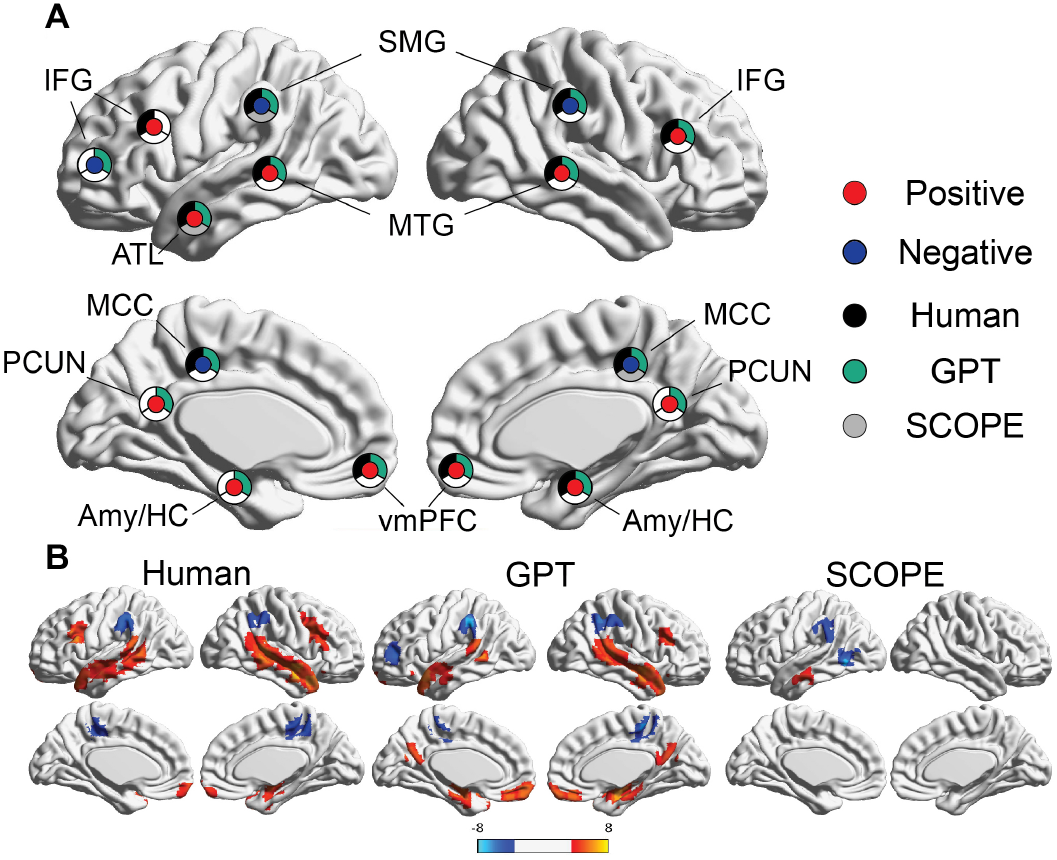
Brain regions identified based on human and GPT valence ratings are remarkably similar. Panel A compared the affective networks shown in Panel B, identified by valence ratings obtained from humans, GPT, and lexical-level (SCOPE) values, after removing the variance explained by arousal and multiple comparisons adjustment (cluster-size-based correction, voxel-wise *p* <.001, cluster size >22 voxels).

**TABLE I.**
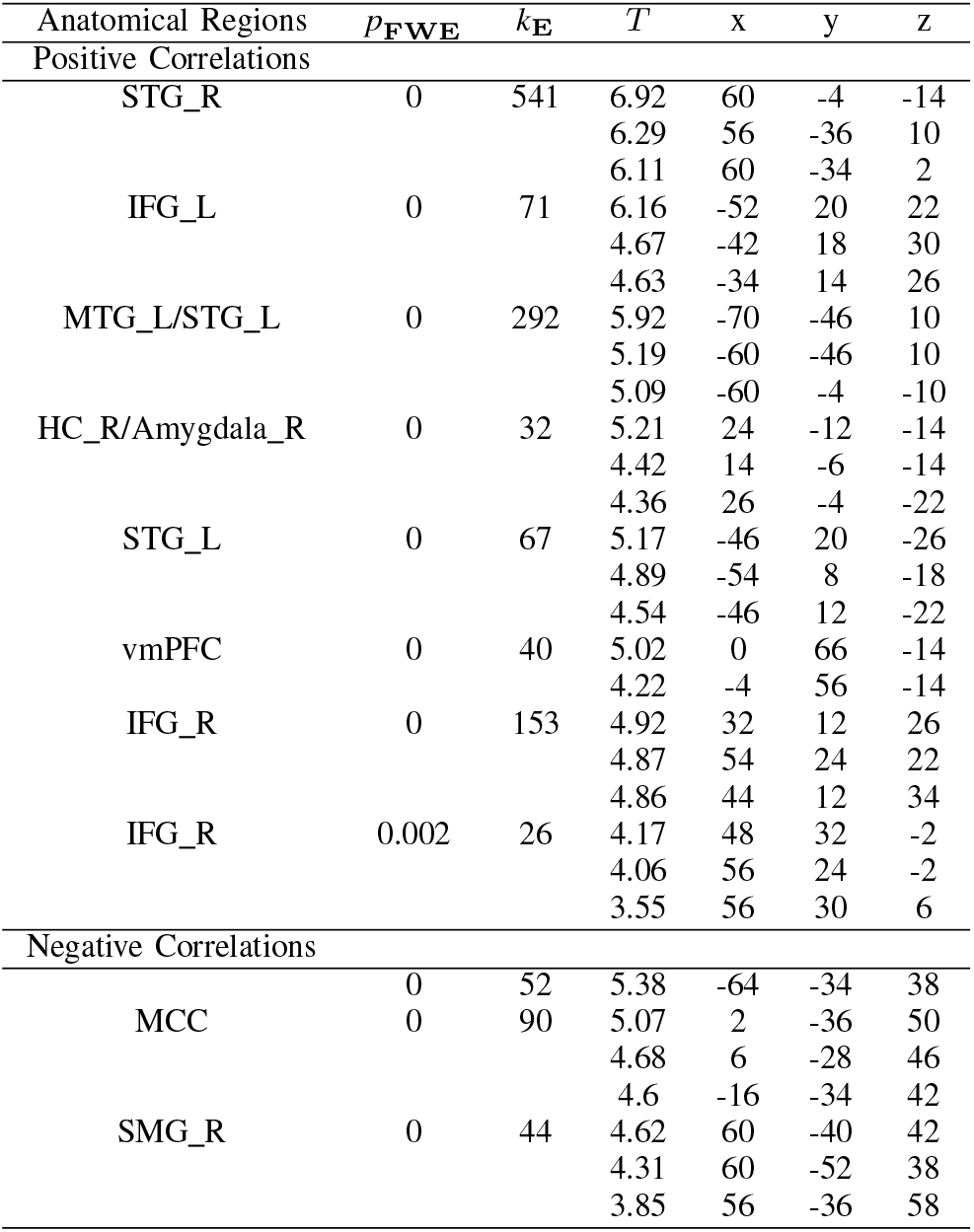
BRAIN REGIONS IDENTIFIED BASED ON HUMAN VALENCE RATINGS.

**TABLE II.** BRAIN REGIONS IDENTIFIED BASED ON GPT VALENCE RATINGS.

| Anatomical Regions | $p_{\text{FWE}}$ | $k_E$ | $T$ | x | y | z |
| --- | --- | --- | --- | --- | --- | --- |
| Positive Correlations |  |  |  |  |  |  |
| MTG_R | 0 | 91 | 6.46 | 30 | -6 | -18 |
| HC_R/Amy_R |  |  | 5.86 | 24 | -12 | -14 |
|  |  |  | 5.01 | 14 | -6 | -14 |
| HC_R/Amy_R | 0 | 78 | 6.19 | -18 | -10 | -14 |
|  |  |  | 5.22 | -30 | -6 | -18 |
|  |  |  | 4.74 | -30 | -4 | -26 |
| MTG_L/STG_L | 0 | 71 | 5.96 | -66 | -46 | 10 |
|  |  |  | 4.82 | -60 | -40 | 18 |
|  |  |  | 3.67 | -66 | -34 | 10 |
| STG_R | 0 | 479 | 5.92 | 60 | -4 | -14 |
|  |  |  | 5.71 | 56 | -36 | 10 |
|  |  |  | 5.46 | 38 | 18 | -34 |
| STG_L | 0 | 127 | 5.75 | -54 | 12 | -14 |
|  |  |  | 5.25 | -46 | 12 | -22 |
|  |  |  | 4.94 | -42 | 20 | -30 |
| vmPFC | 0 | 72 | 5.52 | 0 | 56 | -14 |
|  |  |  | 5.5 | 0 | 42 | -18 |
| PCUN | 0 | 47 | 5.03 | -4 | -64 | 26 |
| IFG_R | 0 | 67 | 4.78 | 42 | 24 | 22 |
|  |  |  | 4.17 | 32 | 12 | 26 |
|  |  |  | 3.78 | 44 | 12 | 34 |
| Negative Correlations |  |  |  |  |  |  |
| SMG_L | 0 | 65 | 6.76 | -64 | -34 | 42 |
|  |  |  | 3.97 | -60 | -48 | 42 |
|  |  |  | 3.64 | -52 | -46 | 42 |
| MCC | 0 | 67 | 6.2 | 2 | -34 | 50 |
|  |  |  | 4.82 | 2 | -40 | 58 |
|  |  |  | 4.33 | 8 | -28 | 46 |
| SMG_L/AG_L | 0 | 117 | 5.82 | 56 | -46 | 46 |
|  |  |  | 4.8 | 56 | -40 | 58 |
|  |  |  | 4.77 | 60 | -52 | 38 |
| IFG_L | 0 | 48 | 5.14 | -40 | 42 | 10 |
|  |  |  | 4.06 | -48 | 44 | 10 |
| MFG_R | 0.004 | 22 | 4.01 | 42 | 36 | 42 |
|  |  |  | 3.9 | 36 | 26 | 42 |

**TABLE III.** BRAIN REGIONS IDENTIFIED BASED ON SCOPE VALUES.

| Anatomical Regions | $p_{\text{FWE}}$ | $k_E$ | $T$ | x | y | z |
| --- | --- | --- | --- | --- | --- | --- |
| Positive Correlations |  |  |  |  |  |  |
| MTG_L | 0.000 | 32 | 4.62 | -64 | -16 | -18 |
| STG_L | 0.000 | 45 | 4.39 | -52 | 8 | -22 |
|  |  |  | 4.07 | -46 | 20 | -30 |
| Negative Correlations |  |  |  |  |  |  |
| MTG_L | 0.000 | 73 | 6.88 | -60 | -60 | 2 |
|  |  |  | 3.82 | -46 | -64 | 2 |
| SMG_L | 0.000 | 51 | 4.59 | -66 | -36 | 38 |
|  |  |  | 4.38 | -60 | -42 | 26 |
|  |  |  | 4.11 | -64 | -48 | 42 |
| MCC | 0.001 | 28 | 4.46 | 6 | -22 | 42 |
|  |  |  | 4.40 | 6 | -24 | 50 |
|  |  |  | 3.87 | 0 | -34 | 50 |
| STG_R | 0.001 | 28 | 4.10 | 42 | -24 | 14 |
|  |  |  | 4.02 | 50 | -10 | 6 |
|  |  |  | 3.68 | 42 | -30 | 22 |

## V. DISCUSSION

Our results demonstrate that GPT ratings are a reliable substitute for time-consuming human ratings. As opposed to previous investigations using GPT, we used the same instructions for GPT and behavioral experiments with humans. Therefore, given the complexity of this multifaceted task, it is not surprising that GPT made some alignment errors. Considering that we presented the instructions and the entire narrative together to GPT to preserve the contextual information, the task GPT encountered was more complex than the human task of reading and rating one segment at a time. GPT needed to use the provided segment markers to separate the narratives into segments, export the first and last words of every segment (for sanity check), and then infer the valence and arousal of every segment. Future studies can test if refining the prompting strategy (e.g., rewarding GPT responses or asking GPT to think step-by-step) can improve output accuracy.

Our results also showed that GPT ratings outperformed the normative affective ratings from single words and were more representative than single human raters, or even the average ratings of a few humans (up to about five). Remarkably, using the GPT-4-1106-preview model, we obtained affective ratings for four narratives in about two hours and at a $5 cost (10 repetitions per narrative). This method was substantially more time- and cost-effective than gathering the ratings from human raters (approximately $2000 for incentive compensation for data collection on Prolific (www.prolific.com), using 40 raters per narrative). This approach is also arguably more accessible for researchers interested in affective sciences who do not have experience with running behavioral studies. Furthermore, GPT is highly accessible to any researchers, as the model has already been trained and does not require advanced programming knowledge. The vision model also opens new avenues for future studies to extract behavioral features from naturalistic materials using the visual modality, such as images and videos. Therefore, the reliability and practicality of using GPT make it suited for automating the feature extraction process for naturalistic materials and implementing ecologically valid research paradigms in both behavioral studies and neuroscience. Besides extracting behavioral ratings from naturalistic stimuli, in combination with automated transcription of audio recordings, GPT could also be used to automate ratings of naturalistic participant responses. Further, our findings on GPT rating of valence and arousal have broader implications and build a foundation to adjust the affective content of material delivered to users in various contexts, such as conversational agents, neuroergonomics, adaptation and personalization of web pages, and multimedia research, to name a few.

## Notes

### Competing Interest Statement

The authors have declared no competing interest.

